# kWIP: The *k*-mer Weighted Inner Product, a *de novo* Estimator of Genetic Similarity

**DOI:** 10.1101/075481

**Authors:** Kevin D. Murray, Christfried Webers, Cheng Soon Ong, Justin Borevitz, Norman Warthmann

## Abstract

Modern genomics techniques generate overwhelming quantities of data. Extracting population genetic variation demands computationally efficient methods to determine genetic relatedness between individuals or samples in an unbiased manner, preferably *de novo*. The rapid and unbiased estimation of genetic relatedness has the potential to overcome reference genome bias, to detect mix-ups early, and to verify that biological replicates belong to the same genetic lineage before conclusions are drawn using mislabelled, or misidentified samples.

We present the *k*-mer Weighted Inner Product (`kWIP`), an assembly-, and alignment-free estimator of genetic similarity. `kWIP` combines a probabilistic data structure with a novel metric, the weighted inner product (WIP), to efficiently calculate pairwise similarity between sequencing runs from their *k*-mer counts. It produces a distance matrix, which can then be further analysed and visualised. Our method does not require prior knowledge of the underlying genomes and applications include detecting sample identity and mix-up, non-obvious genomic variation, and population structure.

We show that `kWIP` can reconstruct the true relatedness between samples from simulated populations. By re-analysing several published datasets we show that our results are consistent with marker-based analyses. `kWIP` is written in C++, licensed under the GNU GPL, and is available from https://github.com/kdmurray91/kwip.

**Author Summary:** Current analysis of the genetic similarity of samples is overly dependent on alignment to reference genomes, which are often unavailable and in any case can introduce bias. We address this limitation by implementing an efficient alignment free sequence comparison algorithm (`kWIP`). The fast, unbiased analysis `kWIP` performs should be conducted in preliminary stages of any analysis to verify experimental designs and sample metadata, catching catastrophic errors earlier.

`kWIP` extends alignment-free sequence comparison methods by operating directly on sequencing reads. `kWIP` uses an entropy-weighted inner product over *k*-mers as a estimator of genetic relatedness. We validate `kWIP` using rigorous simulation experiments. We also demonstrate high sensitivity and accuracy even where there is modest divergence between genomes, and/or when sequencing coverage is low. We show high sensitivity in replicate detection, and faithfully reproduce published reports of population structure and stratification of microbiomes. We provide a reproducible workflow for replicating our validation experiments.

`kWIP` is an efficient, open source software package. Our software is well documented and cross platform, and tutorial-style workflows are provided for new users.

## Introduction

A major application of DNA sequencing is comparing the genetic make-up of samples with one another to either identify commonalities, and thus detect relatedness, or to leverage the differences to elucidate function. Initially, one seeks to confirm assumed genetic lineages and replicates, or to group samples into families, populations, and species. Estimating the genetic relatedness between a broad collection of samples must avoid bias and have minimal per sample cost.

Nowadays, the vast majority of studies in population genomics are performed using next generation sequencing (NGS) [1]. The methods commonly employed to analyse whole genome DNA sequencing data rely on two complementary concepts: the assembly of reference genomes, and comparing samples to this reference by re-sequencing, read mapping, and variant calling. This approach, while functional in model organisms, is not ideal. Selecting the reference individual is mostly random, generating a reference genome assembly is time consuming and costly [2, 3], and analyses based on read alignment to a possibly inappropriate reference genome sequence are highly susceptible to bias [4, 5], to the point where large parts of the genomes are missed when sufficiently different or absent from the reference. Alignment-free methods for measuring genetic relatedness would help overcome this reference genome bias.

Another issue of concern is sample identification. A recent review [6] found that sample misidentification occurs at an alarming rate. With ever increasing sample numbers in (population) genetic projects, the issue of correct and consistent metadata arises on several levels, technical (mix-up) and biological (misidentification). Large field, and entire gene bank collections are being DNA-sequenced. With sample handling from the field through the laboratory to the sequence read files and eventual upload to data repositories, there is ample opportunity for mix-ups and mislabeling of samples and files. This problem is exacerbated by the often highly collaborative nature of such undertakings. Some misidentifications might be virtually undetectable without molecular genetic analysis, such as varying levels of ploidy, cryptic species, or sub-genomes in (compilo)species complexes [7]. Unfortunately, much of this hidden variation is easily overlooked by following aforementioned current best practices to calculate genome-wide genetic relatedness from short read sequencing data. Erroneous sample identification and/or underestimating the level of divergence has implications for downstream analysis choices, such as which samples and populations to use for a Genome Wide Association Study (GWAS); the missing heritability might then in fact be in the metadata.

The field of alignment-free sequence comparison aims to combat these difficulties by avoiding the process of sequence alignment. Approaches include decomposition to words (or *k*-mers) [8–11], sub-string or text processing algorithms [12, 13], and information theoretic measures of sequence similarity or complexity [14]. While avoiding sequence alignment, many alignment-free sequence comparison tools still require prior knowledge of the underlying genome sequences, which precludes their use as a *de novo* tool.

Recently, several algorithms enabling *de novo* comparisons have been published. These extensions all attempt to reconstruct phylogenetic relationships from sequencing reads. `Spaced` [8, 13] uses the Jensen-Shannon distance on spaced seeds (small *k*-mers a short distance from one another or with interspersed disregarded bases) to improve performance of phylogenetic reconstruction. `Cnidaria` [15] and `AAF` [16] use the Jaccard distance to reconstruct phylogenies, while mash [17] uses a MinHash approximation of Jaccard distance to the same effect. None of these methods report acceptable performance within species, meaning their utility in population genomics may be limited.

Here we present a new metric to estimate genetic relatedness that introduces two concepts to *k*-mer-based sequence comparison. Firstly, we no longer compare every *k*-mer, but rather hash them first into a probabilistic data structure. Secondly, we introduce an information theoretic weighting to elevate the relevant genetic signal above the noise. Pairwise similarity is calculated by the inner product between *k*-mer counts, weighted by the information content derived from their population frequencies. Our procedure is implemented in a software tool (`kWIP`). It estimates genetic relatedness directly from sequencing reads by comparing the *k*-mer contents of sequencing runs. We show by simulations and by re-analysing published datasets, that `kWIP` can quickly, and accurately detect genetic relatedness between samples.

## Design and Implementation

`kWIP` operates on files containing sequencing reads generated by common modern sequencing platforms (e.g., Illumina). First, `kWIP` utilises `khmer` [18, 19] to count overlapping words of length *k* (*k*-mers) into a probabilistic data structure (a sketch) for each sample. `kWIP` then counts presence/absence of each *k*-mer across all sample sketches and records this population frequency in a population frequency sketch. We calculate similarity as the inner product between each pair of sample sketches, weighted by the Shannon entropy of the respective frequency. This process is outlined in Fig 1.

**Fig 1.**
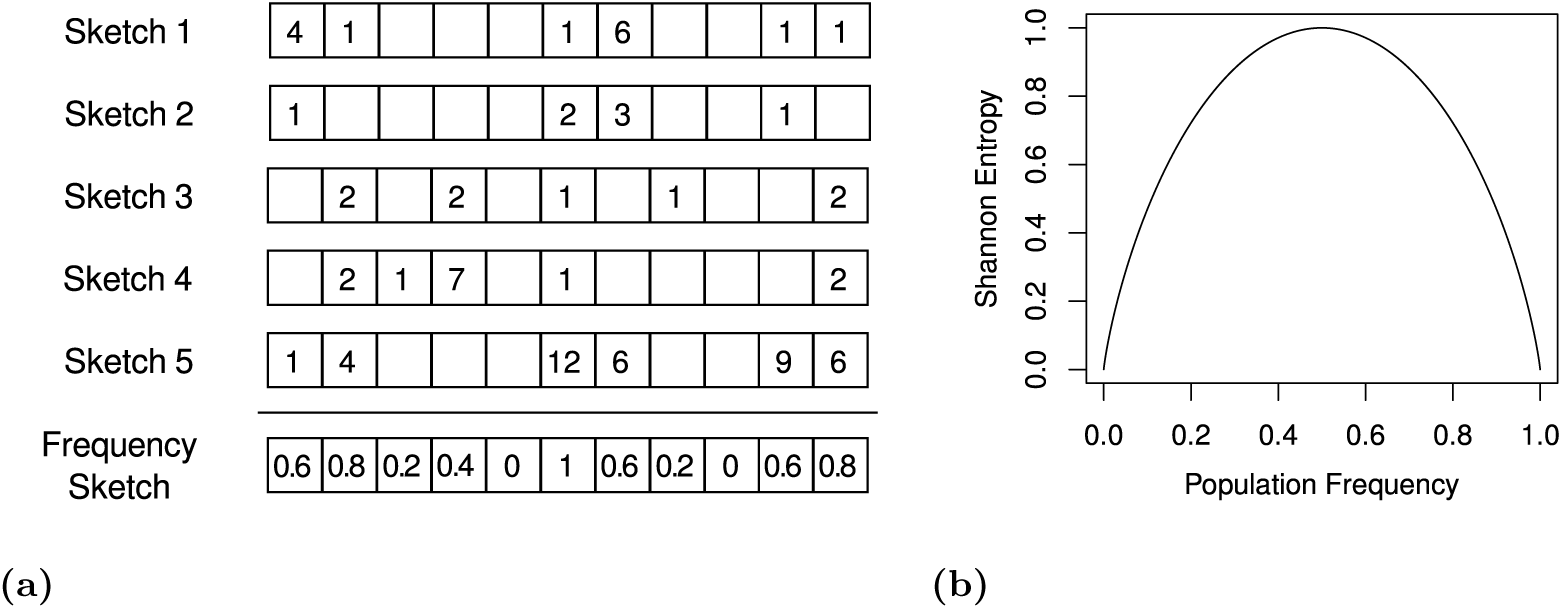
Overview of the Weighted Inner Product metric as implemented in `kWIP`. *k*-mers are counted into sketches (a). The frequencies of non-zero counts across a set of sketches is computed, forming the population frequency sketch. We calculate Shannon entropy of this population frequency sketch as the weight vector for the WIP metric (see equation 2).

### *k*-mer counting

For each sample, `kWIP` uses `khmer` to decompose sequencing reads into overlapping words of some fixed length, e.g., 20. The value of a reversible hash function is computed for each *k*-mer. *k*-mers are canonicalised by using the lexicographically smaller of a *k*-mer and its reverse complement. *k*-mers are counted using one sketch per sample. These sketches are vectors with prime number length, typically several billion elements in size (denoted *S*_*i*_ for sample *i*). The elements of these sketches are referred to as bins (indexed by *b*, e.g. *S*_*ib*_), and can store values between 0 and 255 (integer overflow is prevented). To count a *k*-mer, the *b*-th bin of the sketch (*S*_*ib*_) is incremented, where *b* is the hash value of the *k*-mer modulo the (prime) length of the sketch. For most use cases, we recommend counting *k*-mers between 19 and 21 bases long, as this balances the number of distinct *k*-mers with the uniqueness of each *k*-mer across samples.

Note that the possible number of *k*-mers (4^*k*^) is much larger than the length of a sketch. Therefore, aliasing (or “collisions”) between *k*-mers can occur, but in practice can be avoided with appropriate parameter selection [18]. It is worth noting that aliasing can only increase similarity between any two samples, and this should occur uniformly across all sample pairs.

### Weighting and similarity estimation

Genetic similarity is estimated as the inner product between each pair of sample sketches (*S*_*i*_, *S*_*j*_), weighted by the informational content of each bin. The population sketch (*P*) contains the frequency of occurrence of each bin, calculated as the number of samples with a non-zero count for each bin. We calculate a weight vector (*H*) of these occurrence frequencies using Shannon’s entropy as per Equation (1). In the Weighted Inner Product (WIP) metric (or kernel), similarity is then calculated as the inner product over each sample’s sketch, weighted by *H* as per Equation (2). The unweighted Inner Product (IP) metric is simply the inner product between to sample sketch vectors, *S*_*i*_ · *S*_*j*_. This produces a matrix of pairwise inner products *K*, commonly referred to as a kernel matrix. The kernel matrix is then normalised using the Euclidean norm (3), and converted to distances using the “kernel trick” [20] as per Equation (4). To ensure distance matrices are Euclidean, `kWIP` confirms that the resulting kernel matrix is positive semi-definite by checking that all eigenvalues are non-negative using the Eigen3 library [21].

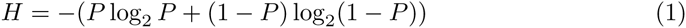

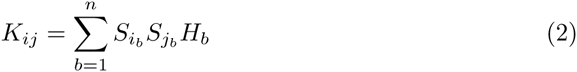

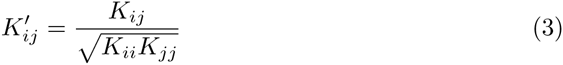

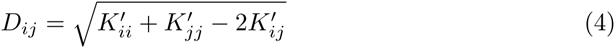

### Implementation

Pairwise calculation of genetic distances from *k*-mer count files with both the WIP and IP metrics is implemented in C++ as `kWIP`. `kWIP` is licensed under the GNU GPL, and source code and pre-compiled executables are available from https://github.com/kdmurray91/kwip. Documentation and tutorials are available from https://kwip.readthedocs.io. To use `kWIP`, one first counts *k*-mers present in each sample using `khmer`’s `load-into-counting.py` script. `kWIP` will then estimate similarity from these counts, producing a normalised Euclidean distance matrix and, optionally, a similarity matrix. `kWIP` parallelises pairwise similarity calculations across cores of a multi-threaded computer to ensure fast operation. Estimating relatedness between 96 rice samples took approximately 4 hours on a 64GB, 16 core (2.6 GHz Sandy Bridge) computer.

## Results

We show that `kWIP` is able to accurately determine genetic relatedness in many scenarios. Using a simulated population re-sequencing experiment, we quantify how the population frequency-based weighting applied by `kWIP` improves accuracy, that is the correlation with the known truth. We recover known technical and biological relationships between sequencing runs of the 3000 Rice Genomes project [22, 23]. We show that a visualisation of `kWIP`’s estimate of genetic relationships between *Chlamydomonas* samples is nearly identical to a similar representation published by the dataset’s authors [24], who used traditional read mapping and variant calling to a reference genome. By analysing a dataset on root-associated microbiomes [25], we show that our approach of sample clustering can be extended to clustering of metagenome samples. See Materials and Methods for detailed description of simulation experiments and validation datasets.

### Quantification of kWIP performance

With simulated data we quantified the performance of `kWIP` comparing our novel metric, the weighted inner product (WIP), with the unweighted inner product (IP), which we consider equivalent to the *D*_2_ statistic. Unsurprisingly, the accuracy, i.e., `kWIP`’s correlation to known truth, decreases with decreasing average sample sequencing depth or genome coverage (Fig 2a); this is true for WIP as well as IP. Importantly, the WIP metric performs far better than IP at low coverages, i.e., below 30-fold. Above a certain coverage, in the case of our simulations at about 50-fold, the performances of the WIP and IP metrics converge. We find that the coefficient of variation between the number of sequencing reads does matter. As a general rule, if a sample has much lower genome coverage than the average, `kWIP` has difficulty accurately determining its relatedness to other samples. Therefore, we advise to exclude such samples or subsample reads from the remainder, if the dataset allows. At constant genome coverage, the improvement in accuracy of the WIP metric relative to the IP metric increases as mean pairwise genetic variation decreases (Fig 2b). While the accuracy of the IP metric decreases markedly below a mean pairwise distance (π) of approximately 0.01, the WIP metric does not show such decrease (Fig 2b).

**Fig 2.**
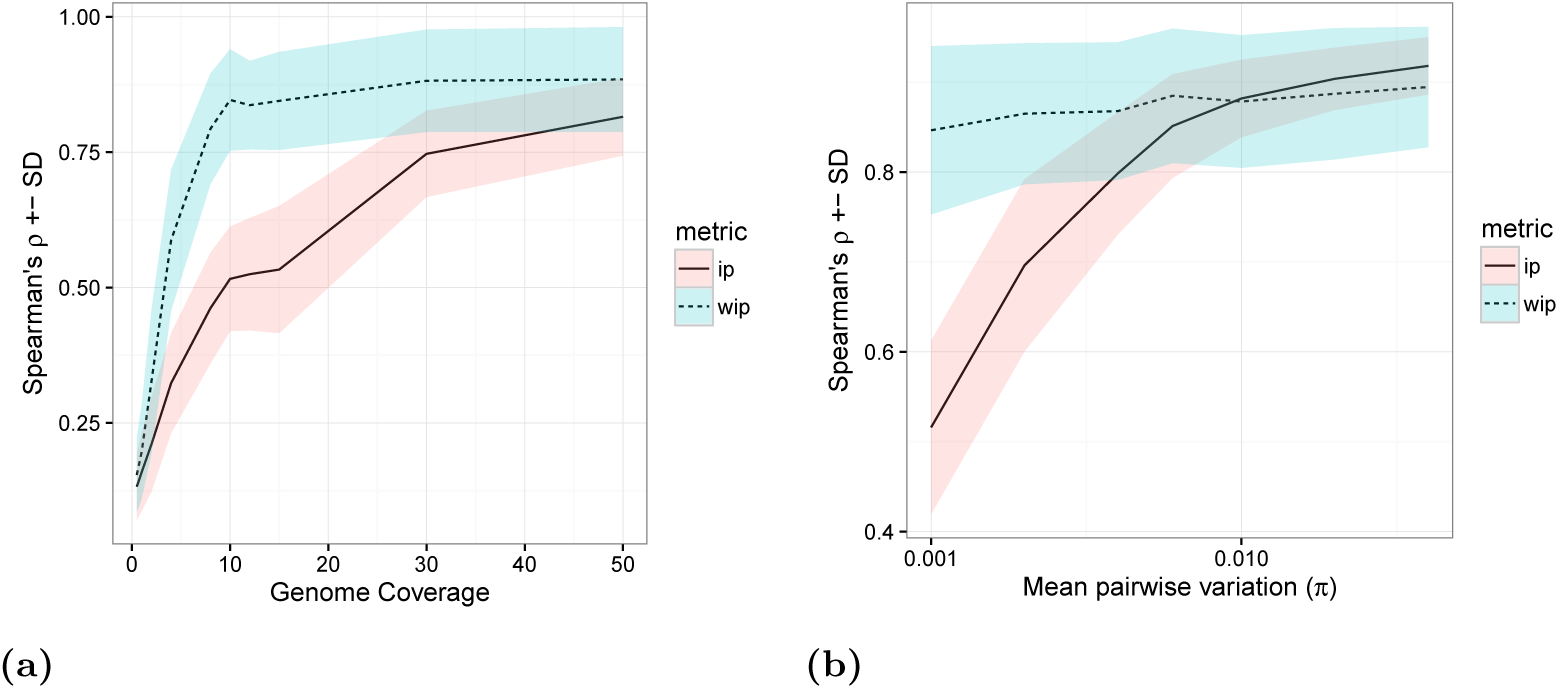
The effect of (a) average sample coverage and (b) average pairwise genetic distance (π) on genetic similarity estimate accuracy. (a) low to moderate coverage (2-30x) weighting increases accuracy, and the weighted metric obtains near-optimal accuracy at 10x coverage compared to above 30x for the unweighted metric. (b) the performance of the unweighted metric decreases rapidly as the mean pairwise distance (π) between samples decreases, however this does not occur for the weighted metric (WIP). The shadings indicate mean ± standard deviation of Spearman’s *ρ* across 50 replicate runs.

### Replicate clustering

`kWIP` can efficiently verify replicates. Figs 3a and 3b show a representative example of replicate clustering. The WIP metric is able to accurately cluster replicates (Fig 3a), whereas the IP metric makes mistakes, as highlighted in red in Fig 3b. We quantified this difference in performance and Fig 3c shows the distribution of rank correlation coefficients between distances obtained with the WIP and IP metrics and the expected clustering patterns for 100 subsets of 96 sequencing runs. The WIP metric outperforms the IP metric, having a higher mean correlation than the IP metric (*p <<* 0.001, paired Student’s T test, *n* = 50).

**Fig 3.**
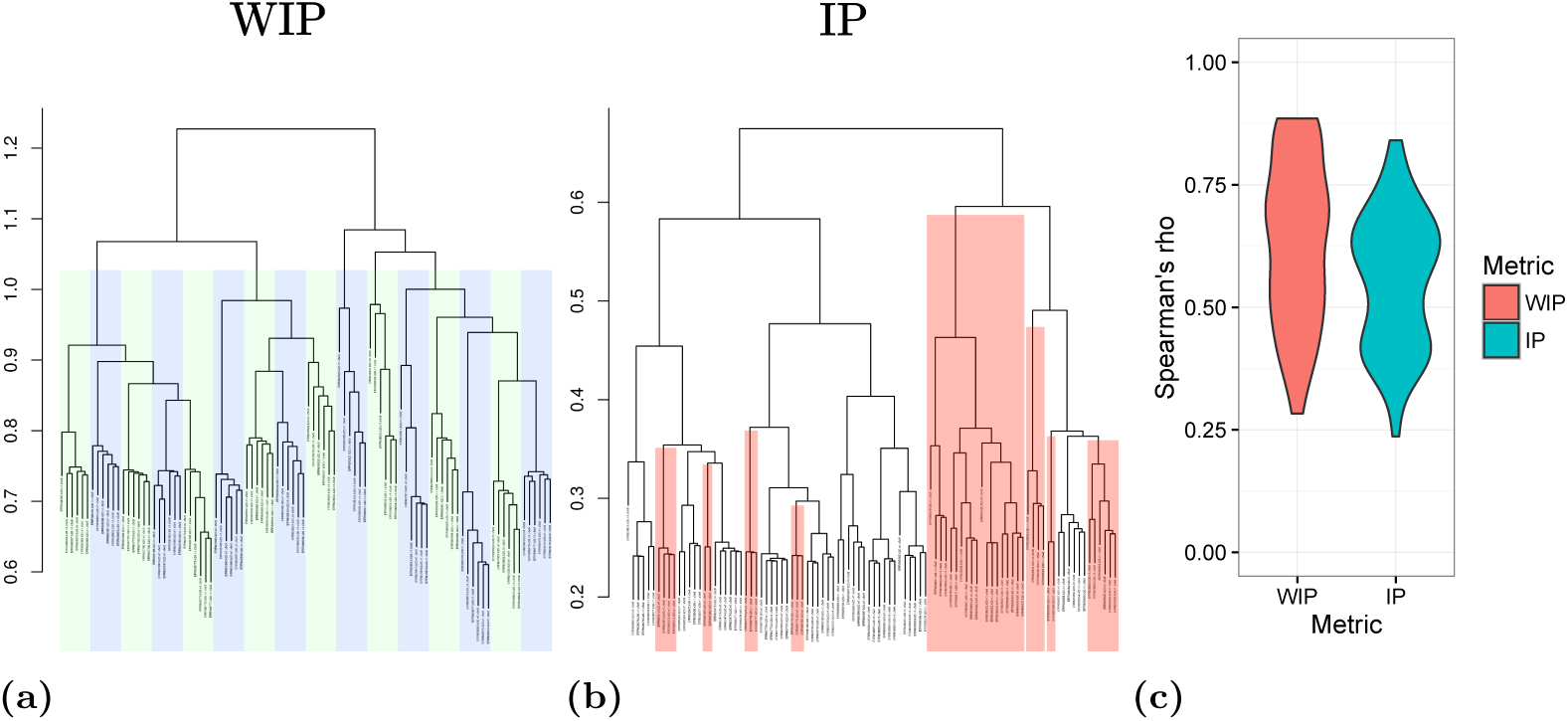
Weighting improves replicate clustering accuracy. (a) and (b) show a representative example demonstrating that the WIP metric (a) correctly clusters all sets of 6 replicate runs into their respective samples (indicated by blue and green bars) while the unweighted metric (b) cannot do so in all cases (indicated by red highlighting). (c) shows rank correlation coefficients to expected relationships over 100 sets of 96 rice runs for the WIP and IP metrics.

### Population structure

Flowers, et al. [24] sequenced 20 strains of *Chlamydomonas reinhardtii* from continental USA, and, by alignment- and SNP-based analysis, find significant population structure, mostly explained by geography [24]. In Fig 4a we display the published genetic relationships as a principal component analysis (PCA) of SNP genotypes exactly as presented by the authors [24]. PC1 separates the laboratory strains (and one western sample) from both eastern and western samples with further structure among wild *Chlamydomonas* collected in western, southeastern and northeastern USA. In Fig 4b we plot the relatedness between the same samples as revealed by `kWIP`, directly from the raw sequencing reads. We note that the results are highly similar.

**Fig 4.**
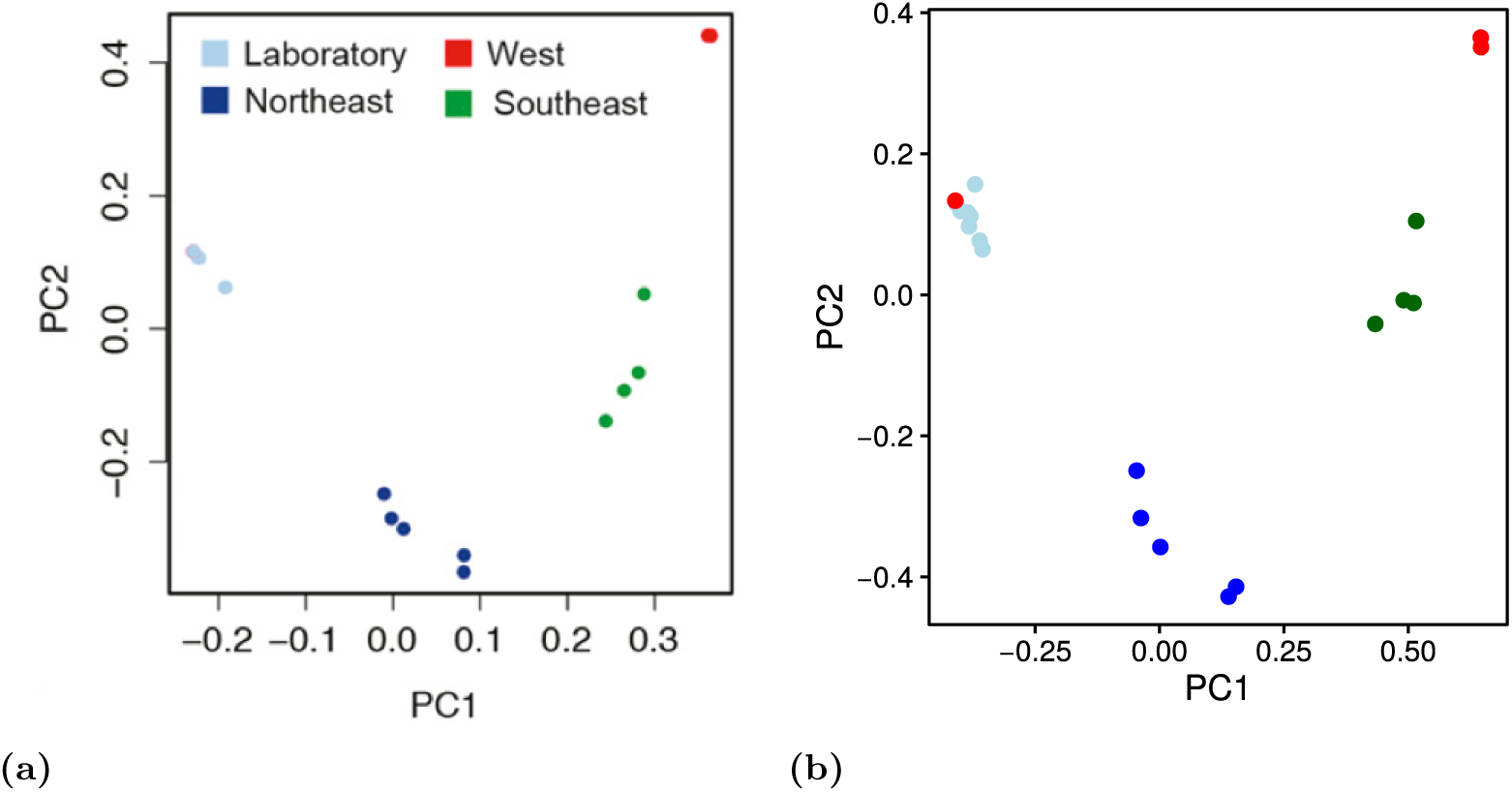
Genetic relatedness between samples of *Chlamydomonas reinhardtii* based on data from [24]. (a) PCA of SNP genotypes reproduced from [24]. (b) Sample relatedness calculated with `kWIP`. Note that in (a), “Sample CC-4414 (red) is obscured behind the cluster of laboratory strains (light blue)” [24].

Each of the 20 strains had been sequenced to a depth of roughly 200-fold genome coverage [24]. By systematically subsampling this dataset we investigate the effect of coverage on the accuracy of `kWIP`’s similarity estimation. Fig 5a shows that as coverage decreases, the accuracy of relationship estimation decreases. We also provide PCA plots of estimated genetic relatedness at varying coverages illustrating this decay 5b. We note that the performance of `kWIP` to determine similarity is very good even at low coverages. A two-fold genome coverage is enough to detect the major splits in this dataset (Laboratory vs West vs East).

**Fig 5.**
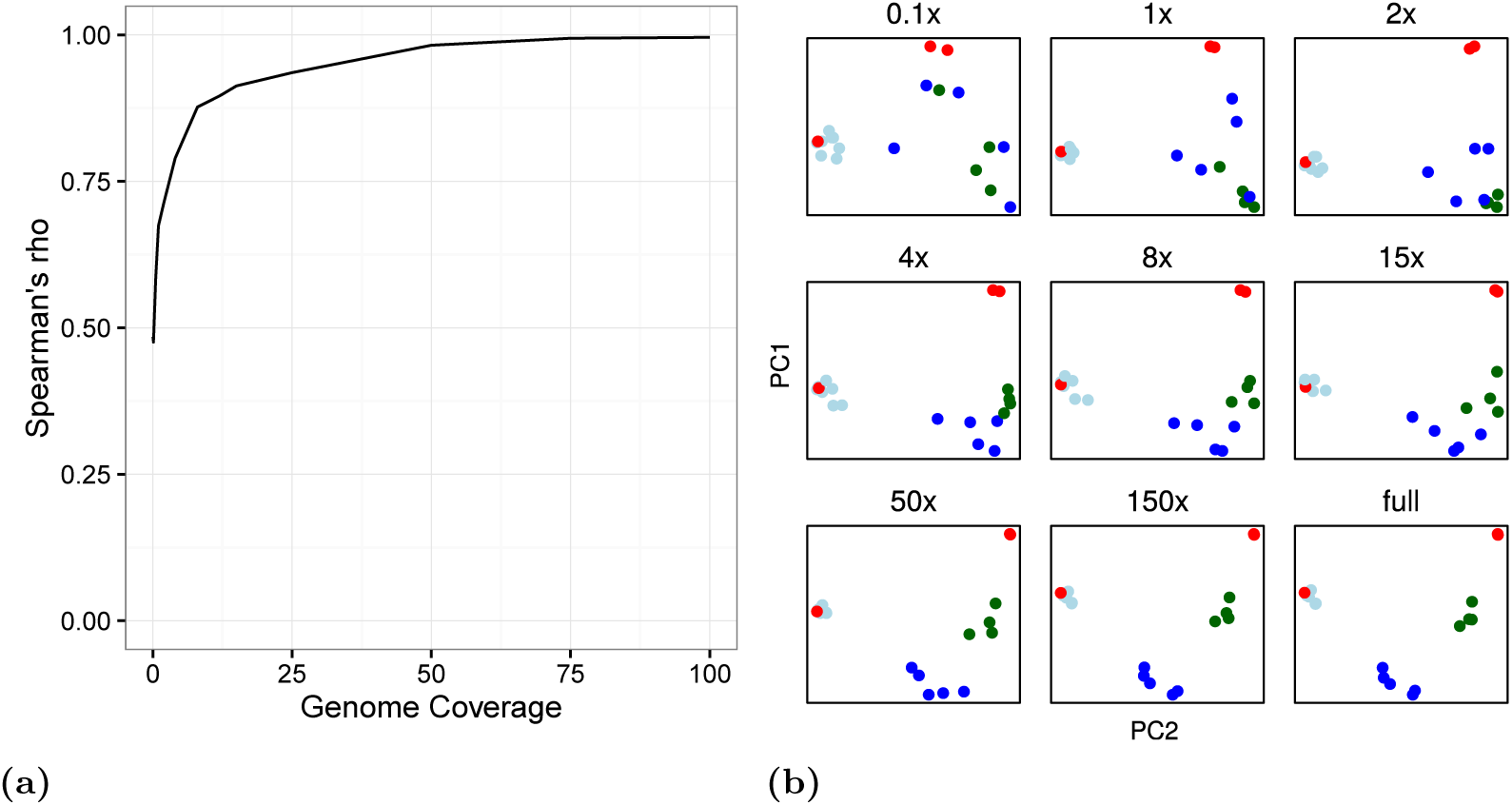
The effect of average sequencing depth (genome coverage) on `kWIP`s estimate of genetic relatedness between samples of *Chlamydomonas reinhardtii* (data from [24]). (a) Spearman’s rank correlation between sub-sampled datasets and the full dataset across a range of subset average genome coverages. (b) PCA plots of relatedness obtained using `kWIP` on selected sub-sampled datasets. “full” refers to the entire dataset (i.e., Fig 4b), while “0.1x” refers to a sub-sampled dataset with average coverage of 0.1 over the *C. reinhardtii* genome (likewise for 1x, 2x, and so on). As noted in 4, a western (red) sample is sometimes obscured behind the cluster of laboratory strains (light blue) [24].

### Metagenome relatedness

Edwards, et al. [25] sequenced rice root-associated microbiomes and find stratification of samples by rhizo-compartment, cultivation site, and cultivation practice. Analysing their raw sequencing data with `kWIP`, we detect highly similar stratification between microbial communities. An example is shown in Fig 6b. We observe a gradient of samples from within the root, through the root-soil interface into soil, and separation by cultivation site. This replicates the separation of samples by rhizo-compartment and cultivation site published by Edwards, et al. [25], shown in Fig 6a.

**Fig 6.**
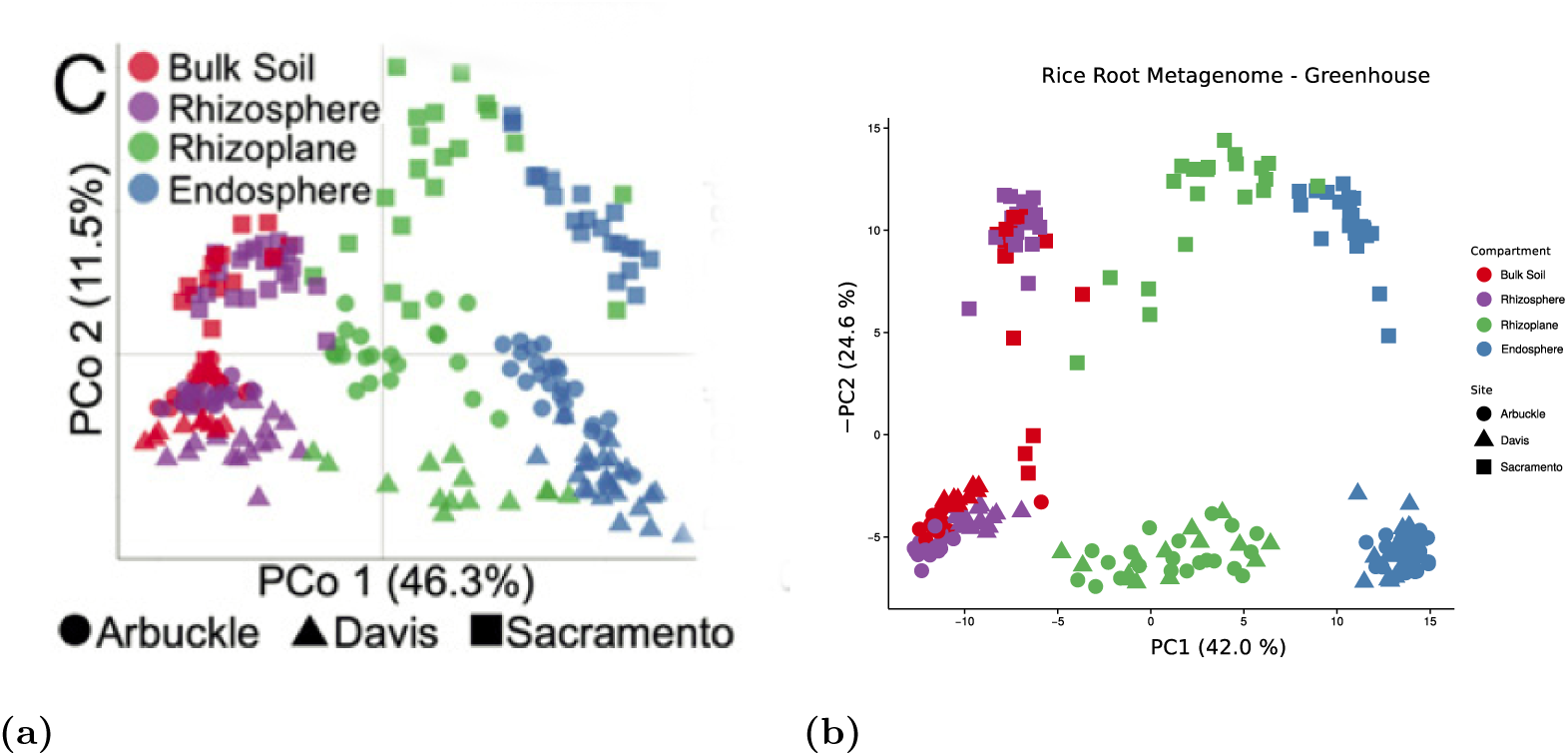
Estimation of similarity between metagenomes. We used `kWIP` to examine the data of [25]. We replicate their observations (a) of stratification of root-associated microbiomes by rhizo-compartment (PC1) and experiment site (PC2). The separation is even more pronounced in the `kWIP` result (b), especially by cultivation site.

## Discussion

The *k*-mer Weighted Inner Product (`kWIP`) estimates genetic distances between samples within a population of samples directly from next generation sequencing data. `kWIP` does not require a reference genome sequence and is able to estimate the genetic distances between samples with less data than is typically used to call SNPs against a reference. As a *k*-mer-based method, `kWIP` is sequencing protocol and platform agnostic, allowing use into the future.

`kWIP` uses a new metric, the weighted inner product (WIP), which aims to reduce the effect of technical and biological noise and elevate the relevant genetic signal by weighting *k*-mer counts by their informational entropy across the analysis set. This weighting has the effect of down-weighting *k*-mers which are either highly abundant, or present in very few samples. Those *k*-mers are typically either common, fixed, repetitive, invariable, or rare, or erroneous. By using Shannon entropy, the weights of common and infrequent *k*-mers are assigned lower, but non-zero weights, allowing some contribution of their signal.

Euclidean distances are then calculated from these weighted inner products and `kWIP` outputs a matrix of pairwise distances between samples, which are easily visualised and may be used for sample classification and to cluster samples into groups. These distance matrices are amenable to quantitative comparison of genetic distance to geographic or environmental distances, for example using mantel tests or generalised dissimilarity modelling. We show high concordance between PCAs obtained using SNP data and those using `kWIP`. It is possible that population genetic statistics, including *F*_*ST*_, could be recovered using `kWIP` via a genealogical interpretation of PCA, as is proposed and shown possible for SNP datasets [26].

We have demonstrated the applicability and effectiveness of `kWIP` using simulations and several published datasets. Through simulations, we quantify how the novel weighting improves accuracy specifically in cases where genetic differentiation or sequencing depth is low (Fig 2a). With data from the 3000 rice genome dataset [23], we reconstruct known relationships between samples and sequencing runs, such as membership of samples to major genetic groups of *Oryza sativa*, and the correct clustering of runs with their replicates (Fig 3). Using a population re-sequencing experiment in *Chlamydomonas* [24] we precisely recreate a visualisation of population relatedness, arguably improving resolution compared to a reference-genome based variant calling approach (Fig 4). This dataset was suitable for comparison because the original authors had based their analysis not only on variants recovered by read alignment against the published reference genome, but attempted to recover and use additional variation by assembling leftover reads that did not match the reference into contigs and calling additional variants between these contigs. This approach, while reducing reference-genome bias, required extensive sequencing depth to enable de-novo assembly; the authors chose around 200-fold coverage, which in turn enabled us to assess `kWIP`’s performance at various sequencing depths (Fig 5).

A current frontier is to efficiently characterise complex metagenome samples. Most studies to date resort to methods of reduced representation. Recently, methods conceptually similar to `kWIP` have been applied to calculate metagenome similarity [17]. We show that `kWIP` is able to detect structure between microbial communities based on 16S rDNA amplicon sequencing data, at least as well as current practice (Fig 6). It should be possible to apply `kWIP` to random shotgun sequencing data from such samples. Besides the similarities between metagenomes, the calculation of intra- and inter-sample diversity is often of interest. The ability to calculate these measures efficiently and *de novo* would be useful because assembling metagenomes accurately is difficult. Estimates of complexity and diversity are currently mostly gene based, but could also be made efficiently at the *k*-mer-level leveraging sketched data structures.

The key innovation of `kWIP` is the combination of a fixed-sized, probabilistic data structure (sketch) for counting *k*-mers with an entropy-weighted inner product as a measure of similarity between samples. By virtue of their fixed size, sketches enable rapid arithmetic operations on *k*-mer counts. Sketches enable `kWIP` to rapidly aggregate across a populations to derive weights, and to efficiently compute the inner products. These benefits outweigh the possibility of collisions between *k*-mers, which in any case have been observed to be rare [18] given appropriate sketch size. Sketching data structures are commonly used for *k*-mer counting (for example Count-Min Sketches [18, 19], and Bloom Filters [27]), but have not been widely adopted in alignment-free sequence comparison.

Weighting of inner products between sketches allows us to account for non-uniform information content of each *k*-mer. `kWIP` weights by Shannon entropy of presence/absence frequency across a population. This provides an assumption-free estimate of the information content of each *k*-mer. By down-weighting both rare *k*-mers introduced by rare variants or sequencing errors, as well as *k*-mers present in most or all samples, we are reducing the contribution of *k*-mers that carry less information. It is possible that other weighting functions that assume various population parameters could provide a more faithful estimate of the information content of each *k*-mer. The application of word-specific weighting has precedence in text processing, where it has been used to account for varying importance of words in a document [28]. However, because we intend `kWIP` to be used in situations where such parameters are either unavailable or potentially inaccurate, we prefer that our weighting is free of assumptions.

An inner product between *k*-mer counts has long been used in the field of alignment-free phylogenetics, where it is referred to as the *D*_2_ statistic. There have been many derivatives of the *D*_2_ statistic that seek to enhance its accuracy where evolutionary distance is large and sites may have mutated multiple times (e.g., 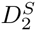, and 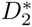 [29–31]). Theortical work has extended the *D*_2_ statistic to calculate phylogenies from NGS data [32, 33]. Use of enhanced models of sequence evolution is not necessary where mutation events occur at independent sites, as is usually the case within populations. `kWIP` does not attempt to re-create evolutionary histories, but rather estimates the similarity of genetic material as it exists today. This is sufficient and even desirable for many of `kWIP`’s intended uses. When validating experimental metadata, one seeks to establish whether similarity between sequencing runs matches expectations. This is particularly true for metagenome samples, where variation can be in both abundance and type of organisms, and their respective genetic sequences. In these cases, estimating present variation between sample genomes, rather than how this variation came to be, is of importance.

Currently, `kWIP`’s strength is estimating genetic similarity between sequencing runs. Because `kWIP` operates reference- and alignment-free, all genetic material present in the sample, the hologenome, will contribute to the analysis. This means that care must be taken that only the organisms of interest have been sequenced. However, we note that *k*-mers that are considered undesirable and chosen to be excluded from the analysis could easily be masked, for example by setting their weight in the weight vector to zero.

Because `kWIP` weights *k*-mers, and hence genome content, based on their frequency in the population being analysed, these weights change when the population changes. This allows for iterative workflows: In a first, all inclusive step the large groupings and outliers are detected; subsequently, subgroups can be analysed with increased resolution.

An important application of `kWIP`, and the WIP metric, is sample classification, where one seeks to compare a sample to some set of known samples. Methods that enable rapid verification of genetic resources, such as stock center accessions, or cell lines, prevent expensive and possibly catastrophic mis-identifications. Sample classification is different from sample clustering as comparisons typically need only be made against a core set of reference samples rather than computing all pair-wise distances between the samples. Inner product kernels have been used to classify protein sequences [34, 35]. To better adapt `kWIP` to classification problems, tree-like structures of kernels [36] or sketches [37, 38] could be explored.

`kWIP` is purposefully designed to operate free of assumptions, or prior knowledge. It is comparing data as presented in the sequencing reads without attempting to reconstruct or approximate the underlying genomes. One could think of several ways of incorporating additional knowledge, which may improve `kWIP`’s power to determine relatedness between underlying genomes. One could, for example, apply ”smoothing” to the *k*-mer counts, with the goal of differentiating between *k*-mers that are genuinely not in the genomes of a sample and those that were not observed due to low coverage and/or stochastic sampling; smoothing is used in natural language modelling [39]. Also, it is possible that alternative distance metrics, such as the Jaccard or Manhattan distances, improve the performance of `kWIP`, which currently uses Euclidian distance. It may further prove valuable to explore spaced seeds [8, 13], or alternative metrics including those considering inexact matches [35, 36].

Estimating the genetic relatedness between a broad collection of natural accessions provides a basis for ecological or functional studies and should be a first step towards solutions in breeding and conservation. In most population level experiments, technical sources of error are dwarfed by the error from insufficient sampling [40]. This is especially true when rare or cryptic lineages are present, and in conditions of non-random mating where population structure is substantial. Such population level noise can only be overcome by broad studies with large numbers of samples, ideally by also merging experiments [41]. When individuals from real-world populations are collected, or collated, there is normally non-uniform genetic relatedness. Initially, one seeks to group samples into more closely related families or more distantly related populations, to then develop core sets for further detailed studies. Genetic outliers occur. They can represent misidentifications and cryptic species and should be excluded. Population re-structuring [40] balances the genetic diversity among a subset of individuals for association studies. *De novo* sample groupings based on whole genome relatedness also inform the selection of suitable reference individuals and/or building the necessary reference genomes. The initial characterisation process must avoid biases and have minimal per sample cost. The use of `kWIP` allows to base the analysis of diversity among samples on low coverage, whole-genome sequence data and thus makes these large, balanced study designs feasible.

More broadly, experiments are condemned to be inconclusive and irreproducible if samples are somehow mislabeled or misidentified. An initial step in all analyses of genetic or functional variation must involve the verification of sample identity [6]. This preliminary analysis should preferably use whole-genome sequence data, be *de novo*, unbiased, and agnostic to sequencing protocol and technology. `kWIP` is an efficient implementation of such a tool.

## Availablity and Future Directions

`kWIP` is implemented in C++ and licensed under the GNU GPL. Source code and pre-compiled executables are available from https://github.com/kdmurray91/kwip. Documentation and tutorials are available from https://kwip.readthedocs.io. The Snakemake workflows and Jupyter notebooks used to perform all analyses presented here are available online at https://github.com/kdmurray91/kwip-experiments; the respective software versions are noted within this repository. Currently, kWIP performs all pairwise comparisons, which scales quadratically (*O*(*n*^2^)) with regards to the number of samples. `kWIP` parallelises pairwise similarity calculations across cores of a multi-threaded computer to ensure fast operation. Analyses of very large data sets, i.e., beyond 10,000s of samples, will benefit from further optimisations to the implementation of `kWIP`, including parallelisation across distributed memory systems with MPI. For each pairwise comparison, the two sketches and the weight vector must fit in main memory. This limits the size of the sketches and the number of pairwise comparisons that will run efficiently in parallel on a given node.

## Materials and Methods

We demonstrate `kWIP`’s performance with both real and simulated datasets. With simulations we quantify the performance of `kWIP`. To demonstrate the utility of `kWIP` in real-world, low-coverage, large-scale population genomics datasets, we analyse data from the 3000 Rice Genomes Project [22, 23]. To show that `kWIP` estimates genetic similarity as well as current best practice SNP-based methods, we re-analysed a population genomics study on 20 strains of *Chlamydomonas reinhardtii* [24] with `kWIP` and compare our result to the published results. Lastly, using data from a study on root-associated microbiomes of rice [25], we show that `kWIP` is able to separate microbial communities from 16S rDNA amplicon data at least as well as current best-practice methods in metagenomics.

We provide all information necessary to reproduce our work: the `kWIP` analyses performed here are implemented in Snakemake workflows [42], which describe all steps and software parameters; random seeds have been fixed where necessary. All downstream analyses are available as Jupyter notebooks [43, 44]. Both the Snakemake workflows and Jupyter notebooks are available online at https://github.com/kdmurray91/kwip-experiments; the respective software versions are noted within this repository.

### Simulations

We simulated several datasets to empirically quantify the performance of `kWIP`. Fifty populations with 12 individuals each were simulated using scrm [45]. Branch lengths within each population were normalised such that the mean pairwise genetic distance (π) was equal. Branch lengths were then scaled over a range of π (between 0.001 and 0.1) to test the effect of mean pairwise genetic distance on `kWIP`’s accuracy. Genome sequences were simulated with DAWG2 [46] and from those short read data for three replicate sequencing runs per individual were generated at various mean coverages (between 0.01- and 200-fold) using Mason2 [47]. We attempted to emulate the reality of sequencing experiments by introducing random variation in read numbers between replicate runs (coefficient of variation of 0.3). We then used khmer to count *k*-mers in these simulated sequencing runs and estimated genetic similarity with `kWIP`, using both the weighted (WIP) and unweighted (IP) metrics.

The performance of our metrics was measured relative to the true pairwise distances between the simulated samples. The true distance matrix between samples was calculated from the simulated, aligned sample genomes (which DAWG2 produces) with `scikit-bio`. Sample-wise distances were replicated three times to allow comparison to the distances obtained from the three simulated sequencing runs. Performance was calculated as Spearman’s rank correlation (ρ) between all pairwise distances using `scipy` [48].

### Datasets

With several published datasets we demonstrate the performance and utility of `kWIP` in real-world scenarios. In all cases, sequence data files for sequencing runs were obtained from the NCBI Short Read Archive using `sra-py` [49]. Reads were extracted using the SRA toolkit to FASTQ files. Low base quality regions were removed using sickle [50] in single-end mode. Counting of *k*-mers into count files (sketches) was performed using the `load-into-counting.py` script of `khmer`. Genetic similarity was estimated using `kWIP`, using the WIP and IP metrics.

To assess how well `kWIP` recovers replicate samples and known sample hierarchies at low sequencing coverage, we turned to publicly available sequence data from the 3000 Rice Genomes project [22, 23]. Samples of the 3000 Rice Genomes project had been sequenced on the Illumina HiSeq2000 platform with technical replicates of individual sequencing libraries split between 6 or more sequencing lanes [22, 23]. Furthermore, there is a rather strong subdivision of rice (*Oryza sativa*) into subgroups. We compiled 100 sets of 96 runs, i.e., for each set we chose 16 samples with 6 replicate runs. We ensured that 8 samples each were described by [22] as belonging to the Indica and Japonica subgroups of *O. sativa*. We estimated the genetic similarity between runs in each of these 100 sets with `kWIP`. The true distances between the different runs in the 3000 rice datasets are not known, but a topology and sample hierarchy can be inferred from the metadata. We hence assessed the performance of `kWIP` in accurately clustering replicates and recovering population structure against a mock distance matrix that reflects the expected topology. We created a distance matrix in which each run had a distance of zero to itself, a distance of 1 to each of its technical replicates (i.e., the other sequencing runs belonging to the same sample), a distance of 2 to each run from other samples in the same rice group (Indica or Japonica), and a distance of 4 to each run from a sample belonging to the respective other rice group. We then used `scipy` to calculate Spearman’s rank correlation between this mock matrix and each distance matrix obtained from real data using `kWIP`. A paired Student’s t-test was performed between the estimates of relatedness from the WIP and IP metrics with the `t.test` function in R. We used hierarchcal clustering to visualise these relationships, performed in also R with the `hclust` function.

We use whole genome sequencing data on 20 strains of *Chlamydomonas reinhardtii* [24] to show the ability of `kWIP` to detect more subtle population structure and to examine the effect of sample sequencing depth (coverage) on the accuracy of `kWIP` in a real-world dataset. Genetic relatedness between the 20 *Chlamydomonas reinhardtii* samples from this study was estimated with `kWIP` using the WIP metric. Classic Multi-dimensional Scaling (MDS) of the `kWIP` distance matrix was performed using the cmdscale function in R. We compare our MDS results against the principal component analysis (PCA) of SNP genotypes as reported by [24]. For Euclidean distance matrices, MDS is equivalent to PCA [51].

We then examined the effect of sample sequencing depth (coverage) on the accuracy of `kWIP` by randomly sub-sampling from the sequencing data of each sample. We sub-sampled to coverages of between 0.01- and 200-fold average coverage (0.01, 0.1, 0.5, 1, 2, 4, 8, 12, 15, 25, 50, 75, 100, 150, 200) across samples using the `sample` command of `seqtk` [52]. We attempted to preserve the coefficient of variation in read numbers that existed in the original dataset (0.12) by sampling a random number of reads from the appropriate normal distribution. Spearman’s rank correlation (ρ) was used to compare pairwise distances calculated at each sub-sampled coverage to those from the original dataset.

To demonstrate that `kWIP` can determine the relatedness of samples in a typical metagenomic dataset, we used next generation sequencing data from a study on rice root associated microbiomes [25] representing 16S rDNA amplicons from soil and root samples. Relatedness between samples was estimated using `kWIP` with the WIP metric, and MDS was performed as above.

## Acknowledgements

We thank Sylvain Forêt, Teresa Neeman, Conrad Burden, Gavin Huttley, Ben Kaehler and Cameron Jack for comments and advice on the metrics, algorithms, and experiments reported here. We thank Luisa Teasdale for comments on earlier versions of this manuscript. This project was supported by the Australian Research Council Centre of Excellence in Plant Energy Biology (CE140100008). This research was undertaken with the assistance of resources from the National Computational Infrastructure (NCI), which is supported by the Australian Government.

